# A novel channel invariant architecture for the segmentation of cells and nuclei in multiplexed images using InstanSeg

**DOI:** 10.1101/2024.09.04.611150

**Authors:** Thibaut Goldsborough, Alan O’Callaghan, Fiona Inglis, Léo Leplat, Andrew Filby, Hakan Bilen, Peter Bankhead

## Abstract

The quantitative analysis of bioimaging data increasingly depends on the accurate segmentation of cells and nuclei, a significant challenge for the analysis of high-plex imaging data. Current deep learning-based approaches to segment cells in multiplexed images require reducing the input to a small and fixed number of input channels, discarding imaging information in the process. We present Channel Net, a novel deep learning architecture for generating three-channel representations of multiplexed images irrespective of the number or ordering of imaged biomarkers. When combined with InstanSeg, ChannelNet sets a new benchmark for the segmentation of cells and nuclei on public multiplexed imaging datasets. We provide an open implementation of our method and integrate it in open source software. Our code and models are available on https://github.com/instanseg/instanseg.

## 1 Introduction

Multiplexed imaging techniques enable the capture of diverse biological markers, offering unprecedented insight into the protein distribution and cellular composition of tissue. Recently developed high-plex imaging methods such as CODEX [1], CyCIF [2] and MIBI [3] enable the capture of dozens of biomarkers in separate imaging channels. While fundamental for the spatial study of tissue, the study of multiplexed images has been hindered by a number of computational challenges. One of these is cell segmentation, in which pixels are assigned to individual cells or cell compartments. Accurate segmentation is crucial for a range of downstream tasks, including cell [4] [5] and tissue [6] phenotyping. Consequently, inaccuracies at the initial stage of segmentation can have far-reaching repercussions in subsequent analyses [7].

Despite the pressing need for accurate and generalized cell segmentation methods, the number of computational solutions currently available to biologists is limited. Recent studies applying cell segmentation to multiplexed images have often relied on CellPose [8] and Mesmer [9]. While both deep learning-based models were trained on diverse datasets and reach human-level performance on selected test sets, the segmentation of cells in multiplexed images is typically restricted to using one or two imaging channels [10]. Indeed, Mesmer only accepts a two channel image, containing one nuclear (e.g DAPI) and one cytoplasmic or membranous marker (e.g E-cadherin). Because it is not possible to rely upon a single marker being available to clearly depict the membrane across all cell types, studies often merge multiple markers into a single channel [5], [11], [12], [13] [14] or select a single marker targeting specific cell sub-populations (e.g. CD45) [15]. Both approaches inevitably discard information that might otherwise have been informative for identifying cell boundaries. Alternatively, both CellPose and Mesmer can be retrained from scratch with a larger number of input channels. However, such models are less general, and would need to be retrained to match the specific combination of markers used for any image set.

Other methods have approximated cell segmentation masks by expanding nuclear masks obtained from single-channel images [16], [17]. The histoCAT platform [18] proposes a two step approach consisting of classifying pixels into three classes (nucleus, membrane and background) using Ilastik [19] followed by segmentation using CellProfiler [20]. This approach requires manually retraining a pixel classifier for images with different biomarker compositions, introducing user-to-user variability. Despite the manual intervention, Ilastik’s segmentation based on conventional machine learning has been shown to produce lower segmentation scores on highly variable datasets when compared to deep learning methods [9].

There therefore remains a need for a generalized computational method that can accurately segment nuclei and whole cells in multiplexed images. Here, we introduce ChannelNet, a novel channel invariant deep learning architecture capable of generating a fixed three-channel representation of multiplexed images irrespective of the nature, number and ordering of biomarkers. We merge ChannelNet into the recent InstanSeg [21] method, substantially outperforming previous methods on public datasets in both accuracy and efficiency. We integrate the method within the popular open-source software QuPath [17] to make InstanSeg (+ChannelNet) amenable for use within existing analysis pipelines, and accessible to biologists with no coding experience.

## 2 Methods

Our contributions to cell segmentation methods are in two parts. Firstly, we build a network that takes an arbitrary number of input channels and generates a fixed three-channel representation. Secondly, we extend our existing InstanSeg [21] method to predict both nuclei and whole cell labels simultaneously. We combine both methods, such that the fixed three-channel representation serves as input to the modified InstanSeg to obtain a channel invariant cell and nucleus segmentation algorithm.

### 2.1 ChannelNet: a Channel Invariant Network optimized for the analysis of multiplexed images

Multiplexed imaging techniques capture a number of biological markers in separate imaging channels. The number and ordering of these markers varies greatly across datasets, and represents a challenge to convolution-based deep learning methods, which are currently rigid to the number and ordering of input channels. We define *channel invariant* to mean any method that is not fixed to take a specific number of input channels and is invariant to permutations of imaging channels, without discarding potentially useful information relating to nucleus or cell boundaries. We present a novel channel invariant module, which we name ChannelNet, that can be introduced in front of a deep learning backbone and imparts channel invariant properties to the network.

#### Intuition

We hypothesise that the task of cell segmentation would be straightforward if the input was a noise-free three channel image, where one channel would depict cell nuclei, a second channel the cytoplasm and a third display cell membranes as clear separating lines. In practice, each channel in a multiplexed histology image typically represents varying amounts of information relating to 1) cell nuclei specifically (e.g. DAPI), 2) a biomarker localized within one or more cell compartments (nucleus, cytoplasm, membrane) for specific subpopulations of cells only, and/or 3) extracellular structures. In principle, any channel might be informative for cell segmentation, but in a way that varies across cell populations. We do not know in general to what extent any individual channel will be informative for the identification of any specific cell compartment.

To output an informative three-channel image from an arbitrary set of imaging channels, a capable channel invariant network needs to (1) estimate the subcellular location of individual biomarkers, (2) infer the location of cell boundaries, even when membrane markers are not present for all cell populations and (3) suppress uninformative or noisy channels, particularly those that redundantly convey information that is more clearly represented in other channels. While some of these tasks may be achieved by treating channels in isolation, an optimal network would require information from all channels to accurately process each individual channel.

#### Our Method

We base our channel invariant network on the U-Net [22] architecture, popular for its ability to extract high-level features while preserving high spatial resolution. Inspired by earlier work to design deep learning networks that operate on sets [23], we adopt a novel approach by representing imaging channels as an unordered set of one channel images, thereby achieving channel invariance by construction. For each block in the U-Net architecture, we first treat the channels in isolation to obtain a number of imaging features, followed by an aggregation step where these features are pooled across all channels and redistributed to the subsequent block by concatenation. The full network is illustrated in Fig. 1.

**Fig. 1:**
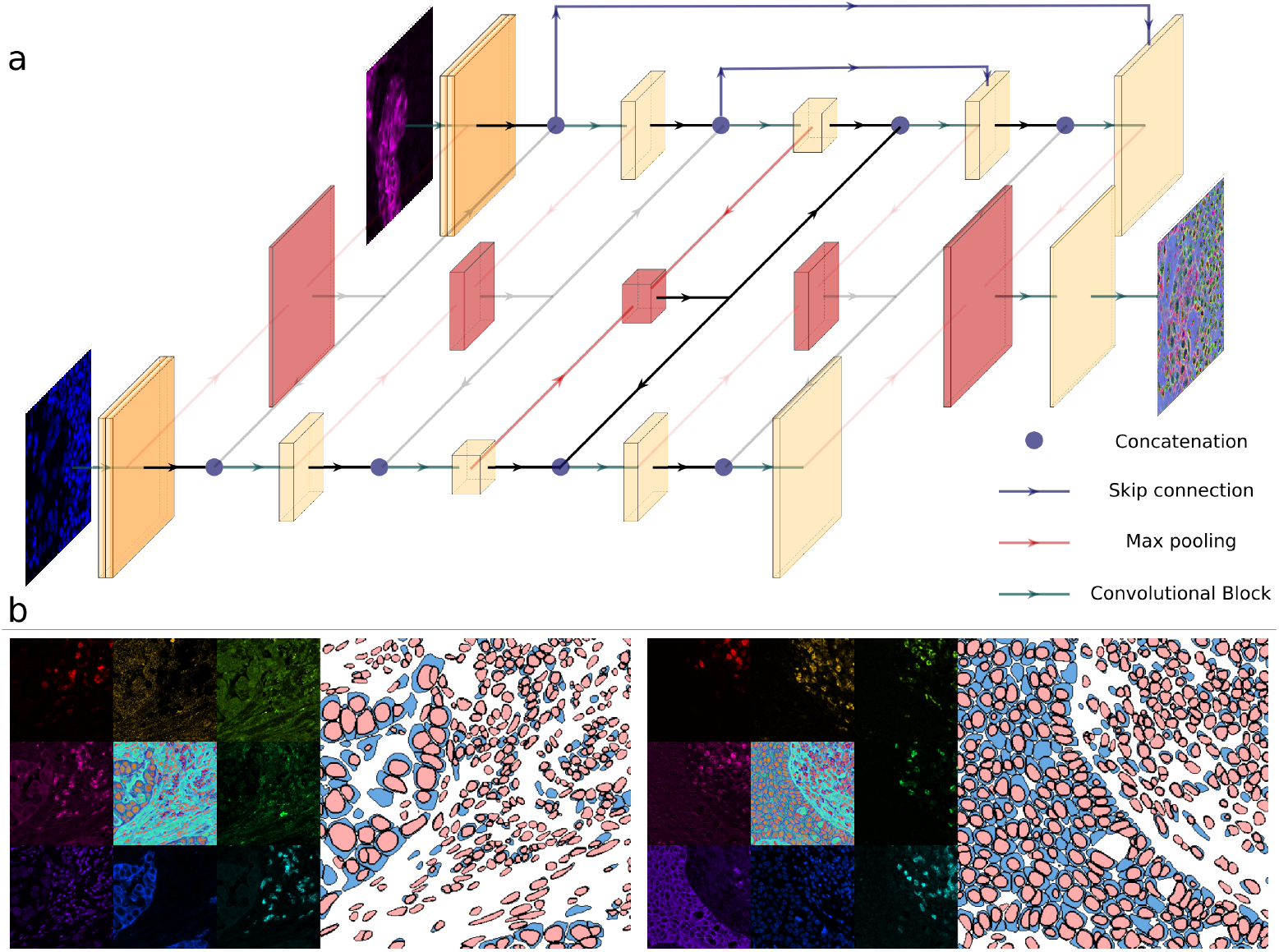
(**a**) The ChannelNet network, based on the U-Net [22] architecture, achieves channel invariance by construction. A multiplexed image is treated by the network as an unordered set of one channel images (only two input channels depicted for simplicity). Identical blocks are used to process each channel in isolation to obtain a number of imaging features (yellow blocks), these are then pooled using a max-pooling operation to obtain shared features (red blocks), that are redistributed to each member of the set via concatenation (blue circles). At the final block, all the imaging features are pooled and a final convolutional block is applied to obtain a fixed three-channel output, which serves as input to the main InstanSeg method for cell and nucleus segmentation. (**b**) Two examples from the CPDMI 2023 dataset. For each, the eight channels are shown in isolation, as well as the three channel ChannelNet output (centre), accompanied by the final nucleus and whole cell segmentation masks predicted by InstanSeg.

In mathematical notation, if we denote *x*_*d*_ the imaging features corresponding to the *d*’th channel, then the *t*’th downsampling block of the ChannelNet architecture performs

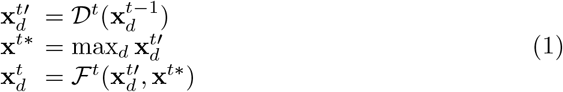

where 𝒟 is a downsampling block with maxpooling and residual connections, and ℱ is a single convolutional block. The *t*’th upsampling block performs:

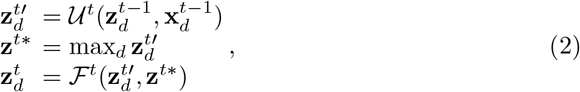

where 𝓊 is an upsampling block with nearest interpolation and residual connections and ℱ is a single convolutional block. Our downsampling and upsampling blocks are inspired from the CellPose backbone architecture blocks [8], also used in the InstanSeg backbone architecture.

### 2.2 Nucleus and whole-cell segmentation using InstanSeg

We extend InstanSeg [21], our recent embedding-based segmentation algorithm, to predict both nuclei and whole cell labels from multiplexed images. To this end, we duplicate the InstanSeg U-Net [22] decoder, so that one branch predicts nucleus labels and the second branch predicts whole cell labels. During training, we only compute and backpropagate the loss using whichever labels are present in the ground truth. This method allows for the simultaneous prediction of nucleus and cell labels even when paired labels are not present in the ground truth.

We treat the ChannelNet network as part of the InstanSeg method, with no additional training steps or loss function required. In effect, ChannelNet is trained to minimize the overall segmentation loss by optimizing the input of the main InstanSeg network. As a result, the three-channel output is not forced to correspond to nuclear, cytoplasmic or membranous signals: any informative representation may be learned, but we expect these to correlate with cell compartments.

### 2.3 Datasets

#### TissueNet

(Modified Apache, Non-Commercial) [9] We use TissueNet v1.1 to benchmark the of segmentation performance of InstanSeg. TissueNet is a large dataset collated from a range of microscopy platforms. The images consist of two channels, one containing a nuclear marker (e.g. DAPI) and a separate containing a single cytoplasmic or membrane marker (e.g. E-cadherin).

#### CPDMI 2023

(CC BY 4.0) [24] The Cross-Platform Dataset of Multiplex fluorescent cellular object Image annotations (CPDMI) is a dataset of multiplexed fluorescence images from various human organs including lung, breast, pancreas, colon, lymph node, ovary, skin, tongue, sacrum, lymph node, hypopharnyx, spleen and tonsil. The images were obtained using the Akoya Vectra, Zeiss Axioscan and Akoya CODEX platforms. Whole cell and/or nucleus annotations are provided for small crops of each of the images based on hand drawn annotations from multiple annotators and reviewed by a pathologist. Altogether, the dataset contains nearly 50 different biomarkers, with individual images containing between 8 and 32 channels. We use this dataset extensively to develop and validate our channel invariant methods. We combine the Vectra and Zeiss images to form our training and validation splits and reserve all the CODEX images to form the test split.

### 2.4 Benchmarks

The number and order of channels differ throughout the CPDMI 2023 dataset. While this is supported by our proposed approach, no previous cell segmentation methods provide channel invariance to enable direct comparison. Consequently, we benchmark our method in two steps: (1) we compare InstanSeg to existing cell and nucleus segmentation methods on an existing fluorescence imaging dataset with fixed number of channels, and (2) we compare our channel aggregation method to previous channel aggregation strategies on an existing fluorescence imaging dataset with a variable number of channels.

For the first step, we benchmark our proposed ChannelNet + InstanSeg nucleus and cell segmentation pipeline against Mesmer [9] on the TissueNet dataset [9], consisting solely of two channel images. We use the public Mesmer model, trained on identical train, validation and test splits. For both Mesmer and InstanSeg, we resize images to 0.5 microns per pixel, as required by the models. Timing was performed on a laptop with a Quadro RTX 3000, 6GB GPU with a batch size of 1.

For the second step, (1) we investigate whether our channel aggregation strategy harms the segmentation accuracy of InstanSeg on the TissueNet dataset, which contains a fixed number of channels. (2) We compare our channel aggregation strategy to other methods that are currently being used to merge channels [5], and (3) investigate how increasing the number of input channels affects the final segmentation accuracy. Finally, (4) we investigate the effect of ablating the ChannelNet architecture on the accuracy of InstanSeg’s predictions.

### 2.5 Channel aggregation baselines

Previous studies applying deep learning-based cell segmentation to multiplexed images have generated two-channel images, with one containing a nucleus marker and a second summing the cytoplasmic markers [5] [11]. On the CPDMI 2023 dataset, we use the ground truth labels to determine the subcellular location of each marker, and use this information to merge markers expressed predominantly in the cell nucleus into one channel and cytoplasmic markers in a separate channel. Specifically, if the mean channel intensity under the nucleus labels was larger than the mean intensity under the whole cell labels, the marker was determined to be expressed in the nucleus and conversely for cytoplasmic markers. Note that we had to look at the test set labels for determining the location of some biomarkers that did not appear in the training split.

We add a third baseline where we generate three-channel (RGB) images by randomly assigning primary and secondary colours to each of the input channels. This RGB projection is often used for visualization purposes and may enable the separation of touching cells visually. In this baseline, we ensure that the channel corresponding to the nuclear marker is always in the same channel. Examples of the three-channel and two-channel channel aggregation baselines are shown in Supplementary Fig. 1.

### 2.6 Ablation

For our ablation study, we prevent ChannelNet from sharing information across the channels. In practice we set **x**^*t*∗^ in Eqn. 1 and **z**^*t*∗^ in Eqn. 2 to zeros. This is equivalent to suppressing all but the last red block in Fig. 1.

For all our ablations and channel aggregation baselines, we train separate models from scratch on identical train, validation and test splits.

### 2.7 Evaluation Metrics

The *F*_1_ score was used as a metric for detection accuracy. We report both 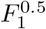, determined at the Intersection over Union (IoU) threshold of *τ* = 0.5 and 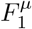, calculated as the mean *F*_1_ score over the interval [0.5, 0.9] with a step of 0.1. We also report Segmentation Quality (*SQ*) defined as the average IoU of all correct matches (above the IoU threshold of *τ* = 0.5). Our metrics are calculated using the Stardist implementation^1^.

### 2.8 Preprocessing

We scale the input image so as to set the 0.1% and 99.9% percentiles of the pixel values to 0 and 1 respectively, independently across channels. We resize images to 0.5 microns per pixel using bilinear interpolation.

### 2.9 Augmentations

For the benchmarks on the TissueNet dataset, we use minimal augmentations during training so as to fairly compare results with other methods. These include horizontal and vertical flips, axis-aligned rotations and random crops of size 256 ×256.

For additional models that we make publicly available, we use further augmentations, including concatenating up to 30 duplicated channels with various amounts of Poisson noise, followed by channel suppression with probability 0.3. We also perform histogram normalization, contrast and brightness shifts.

### 2.10 Training Details

We train ChannelNet and InstanSeg in a single training loop comprising 500 epochs, each consisting of 1000 batches of size 3. To batch images with a variable number of channels, we concatenate extra empty channels to make all batched images the same size. We train on 256 ×256 pixel crops, using the Adam optimizer and a fixed learning rate of 0.001.

### 2.11 Assigning nuclei to cells

Assigning individual nuclei to their respective cells is a non-trivial and often ignored issue in cell segmentation. The difficulty arises when (1) multiple nuclei are predicted within a single cell, (2) a nucleus does not overlap with any predicted whole cell, (3) a nucleus is not fully contained by a corresponding cell or overlaps with multiple cells. As these labeling inconsistencies are present in the training and testing datasets, we do not post-process the segmentation predictions to match individual nuclei and cells. However, we provide the tools to resolve such inconsistencies in our public implementation to facilitate downstream tasks. Based on the observation that nucleus segmentation is typically an easier task compared to whole cell segmentation, our approach, illustrated in Fig. 2, is in two steps. For each cell overlapping with one or more nuclei, we match it to the nucleus with the highest overlap. We then define the cell mask as the union of the initial cell mask and the matched nucleus mask. For any nucleus that remains unmatched, we create a new cell mask that exactly corresponds to the nucleus mask.

**Fig. 2:**
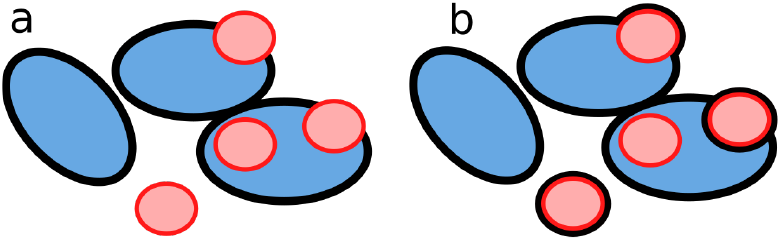
Sketch showing the result of our nuclei to cell assignment steps. Nuclei and cells are predicted independently by InstanSeg (a), and can later be resolved for downstream analysis (b). Black lines: cell boundaries, red lines: nucleus boundaries.

## 3 Results

### 3.1 InstanSeg (+ChannelNet) sets a new state of the art for the segmentation of cells and nuclei in multiplexed images

We report our benchmarking results on the TissueNet test set in Tab. 2. We find that the InstanSeg base model, here accepting only two input channels, outperforms Mesmer for both nucleus and whole cell segmentation. We further show that adding a ChannelNet adaptor, which loses information on the ordering or number of input channels, results in little degradation in the segmentation accuracy.

**Tab. 1:** Total number of images, nucleus and cell annotation counts for each dataset (Tr: Train,V: Validation,T: Test, N: Nuclei, C: Cells)

| Dataset | Channels | Images (Tr/V/T) | Instances (Tr/V/T) |
| --- | --- | --- | --- |
| CPDMI 2023 [24] | 8 - 32 | 98 / 25 / 10 | 15,374 / 4,100 / 1,355 (N)<br>47,162 / 12,139 / 5,880 (C) |
| TissueNet [9] | 2 | 2,580 / 3,118 / 1,324 | 932,591 / 275,495 / 135,633 (N)<br>988,150 / 294,347 / 145,222 (C) |

**Tab. 2:** Quantitative segmentation results on the TissueNet test set. Note that some of the test set labels were curated using a Mesmer model, as described in [9].

| Method | Target | $F_1^\mu$ | $F_1^{0.5}$ | $SQ$ | Time (s) | Images/second |
| --- | --- | --- | --- | --- | --- | --- |
| Mesmer | Nuclei | 0.7115 | 0.9030 | 0.8421 | 280 | 4.7 |
|  | Cells | 0.6328 | 0.8593 | 0.8163 |  |  |
| InstanSeg | Nuclei | **0.7760** | 0.9160 | **0.8738** | **31** | **42.7** |
|  | Cells | **0.6725** | 0.8699 | **0.8343** |  |  |
| InstanSeg (+ ChannelNet) | Nuclei | 0.7649 | **0.9207** | 0.8646 | 36 | 36.8 |
|  | Cells | 0.6654 | **0.8811** | 0.8252 |  |  |

We also show that InstanSeg is a highly efficient cell segmentation algorithm. The processing speed on the TissueNet test set containing 1,324 images of shape 256× 256 pixels was up to 42.7 images per second on a laptop GPU.

### 3.2 InstanSeg (+ChannelNet) allows for the accurate segmentation of cells and nuclei in images with varying number and ordering of channels

We report the segmentation accuracy of InstanSeg on the CPDMI 2023 dataset, comprising images with between 8 and 32 channels, in Tab. 3. InstanSeg predicted nucleus and whole cell labels with an accuracy of 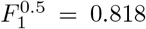 and 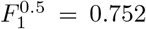 respectively. The segmentation accuracy was substantially higher than the two-channel baseline which separated nuclei and cytoplasmic markers, as used in previous studies [5]. Our three-channel-projection baseline performed worse, despite such RGB projections being the standard for visualizing multiplexed images in software. Compared to the other baselines, our channel aggregation strategy has the advantage of not requiring user input on the subcellular location of the markers. Furthermore, we show that ablating the ChannelNet adaptor degraded segmentation performance, thereby demonstrating the benefit of cross-channel information pooling.

**Tab. 3:** Baseline and ablation study on the CPDMI 2023 validation set.

| Method | Target | $F_1^\mu$ | $F_1^{0.5}$ | $SQ$ |
| --- | --- | --- | --- | --- |
| InstanSeg (+ ChannelNet) | Nuclei | **0.522** | **0.818** | **0.767** |
|  | Cells | **0.438** | **0.752** | **0.741** |
| InstanSeg - two channel (No ChannelNet) | Nuclei | 0.498 | 0.798 | 0.761 |
|  | Cells | 0.380 | 0.710 | 0.716 |
| InstanSeg - three channel (No ChannelNet) | Nuclei | 0.330 | 0.638 | 0.707 |
|  | Cells | 0.345 | 0.662 | 0.709 |
| InstanSeg (+ ablated ChannelNet) | Nuclei | 0.491 | 0.798 | 0.757 |
|  | Cells | 0.418 | 0.738 | 0.732 |

We show qualitative segmentation outputs in Fig. 3. As hypothesized, the fixed three-channel ChannelNet outputs correlated with cell compartments, shown by the strong nuclear, cytoplasmic or membranous signals. We stress that ChannelNet is only trained to minimise the segmentation error of the main InstanSeg network, so intermediate representations may differ upon retraining.

**Fig. 3:**
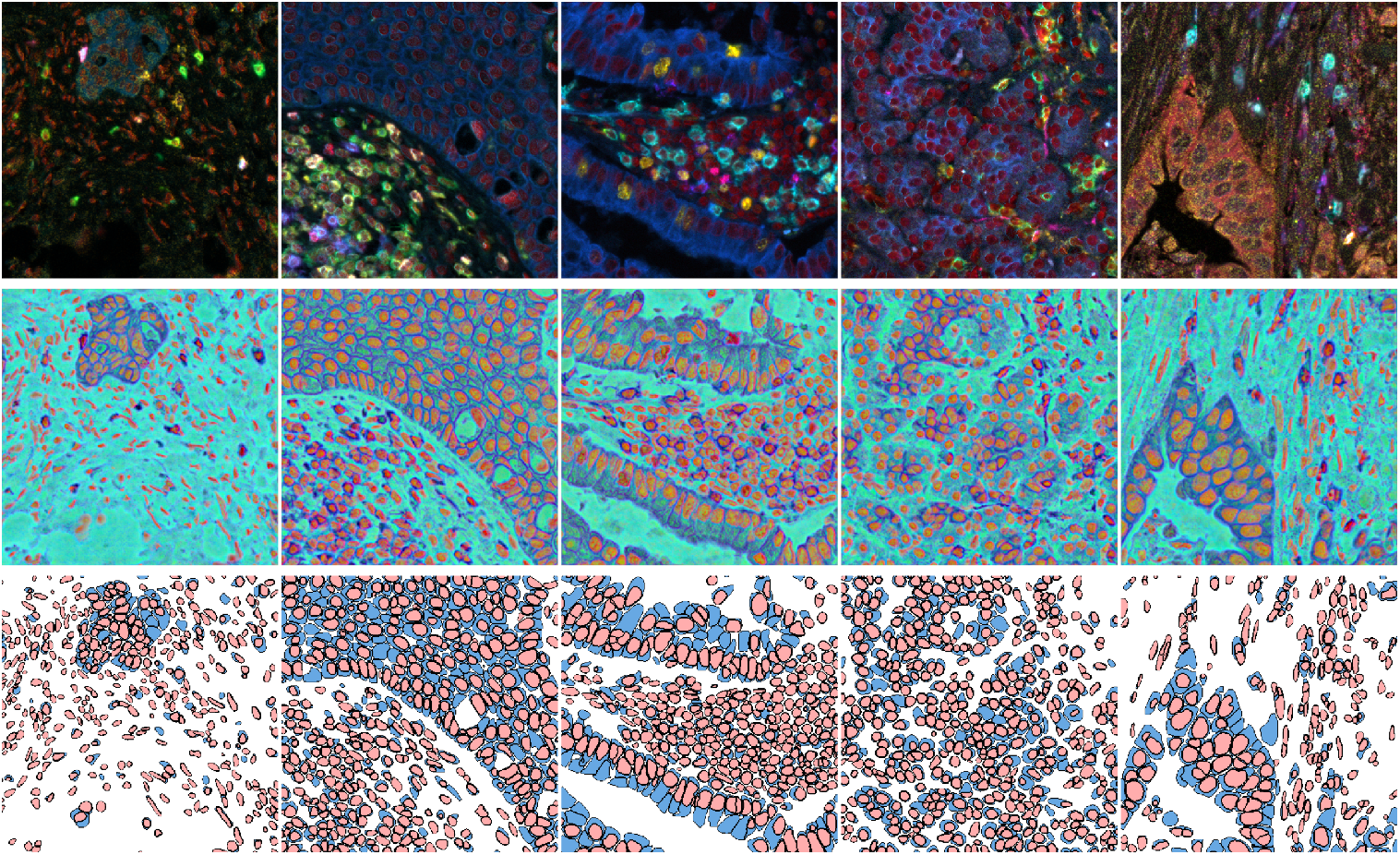
Qualitative results of InstanSeg (+ ChannelNet) on the CPDMI 2023 dataset. Top row: five multiplexed images (rendered in RGB for display). Note that the number and ordering or colour channels is not conserved across experiments. Middle row: corresponding fixed three-channel representation generated by ChannelNet. Note the consistency of the nuclear, cytoplasmic and membranous signals. Bottom row: final nucleus and cell segmentation masks predicted by the InstanSeg (+ChannelNet) method.

#### 3.2.1 InstanSeg (+ChannelNet) segmentation accuracy improves with increasing number of imaging channels

We test InstanSeg (+ChannelNet) on the CPDMI 2023 test set consisting of 28-32 channel CODEX images. We show that the accuracy of the whole cell predictions increase monotonically as the number of input channels was increased. Our method predicted cellular labels with an 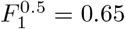when a single DAPI channel was provided, and 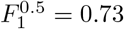 when all channels were provided Fig. 4. We show a similar trend when the *N* most informative input channels were selected. We show qualitatively the effect of increasing the number of input channels from one to seven in Supplementary Fig. 2.

**Fig. 4:**
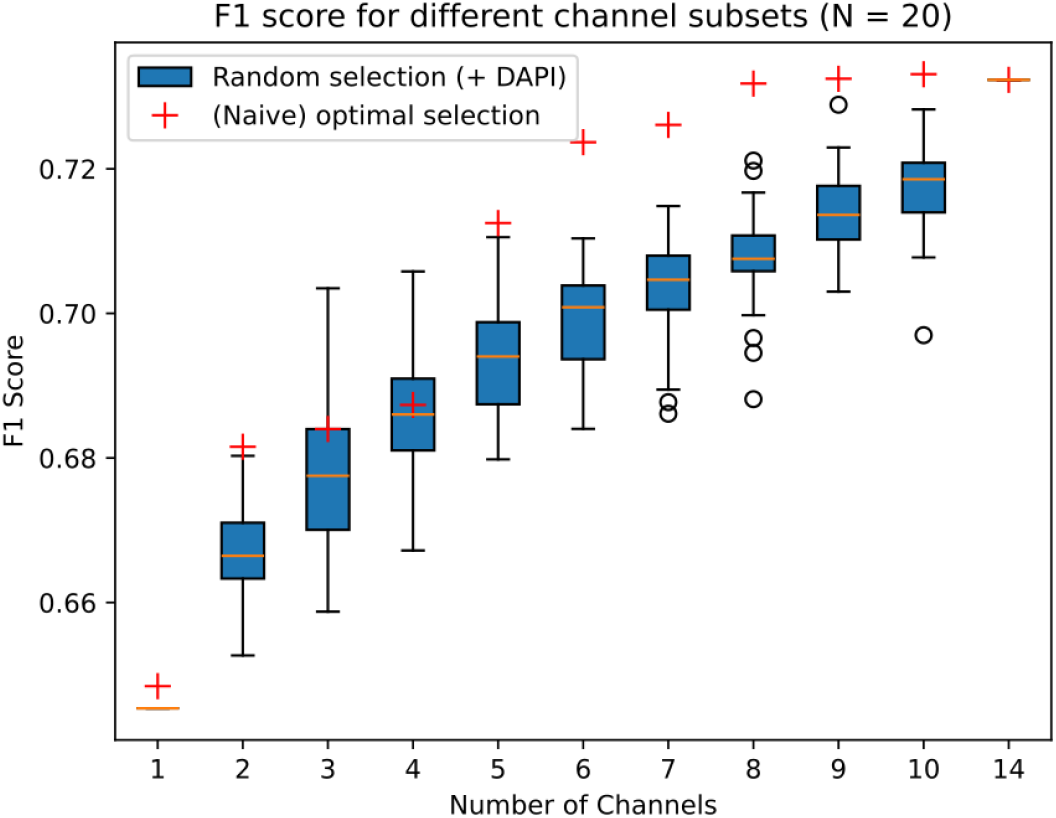
On the CPDMI 2023 test set, consisting of 13-14 non-redundant channel images, we show the F1 score of the whole cell predictions as we increase the number of randomly sampled channels (blue boxes) for N=20 repeats. In each case we ensure that a DAPI channel is selected. Next, we rank each channel by its informativeness based on F1 score when using that channel alone. We then sample the top N most informative channels and evaluate InstanSeg on their combination (red crosses).

#### 3.2.2 InstanSeg enables accurate analysis of multiplexed images

InstanSeg provides segmentation masks for both nuclei and whole cells. We find that this dual prediction enables downstream analyses, such as predicting the subcellular location of biomarkers and nucleus to cell area ratios. We compare InstanSeg’s predictions against ground truth annotations on the CPDMI 2023 dataset in Fig. 5, and show that InstanSeg’s downstream predictions strongly agree with those from the ground truth annotations.

**Fig. 5:**
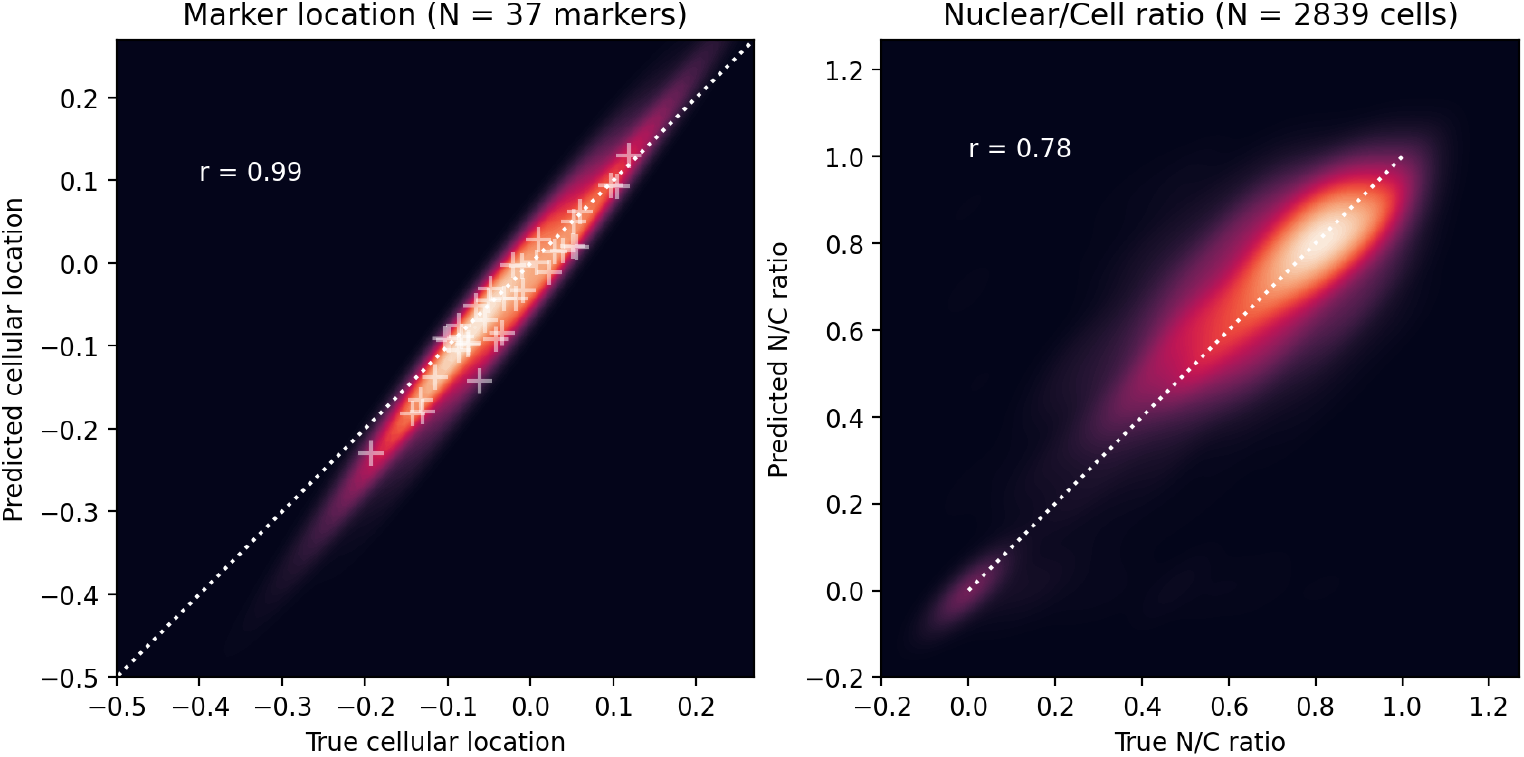
**Left** For each marker in the CPDMI 2023 validation dataset, we compare the predicted cellular location log_2_(nuclear*/*cell) of each marker compared to the true location obtained from the ground truth labels. Note that markers/cells were not gated for this analysis. **Right** Predicted nucleus to cell area ratio (N/C ratio) of InstanSeg predictions versus ground truth labels on the CPDMI 2023 validation dataset

### 3.3 QuPath extension

All aspects of the prediction - including pre- and post-processing - are encapsulated in a single TorchScript file, greatly facilitating the implementation of the method into other software. We have demonstrated this by building a QuPath [17] extension, enabling biologists to use InstanSeg (+ChannelNet) with no coding experience. Our extension supports GPU acceleration on both NVIDIA and Apple hardware, and provides a user friendly interface to select imaging channels for segmentation (Fig. 6).

**Fig. 6:**
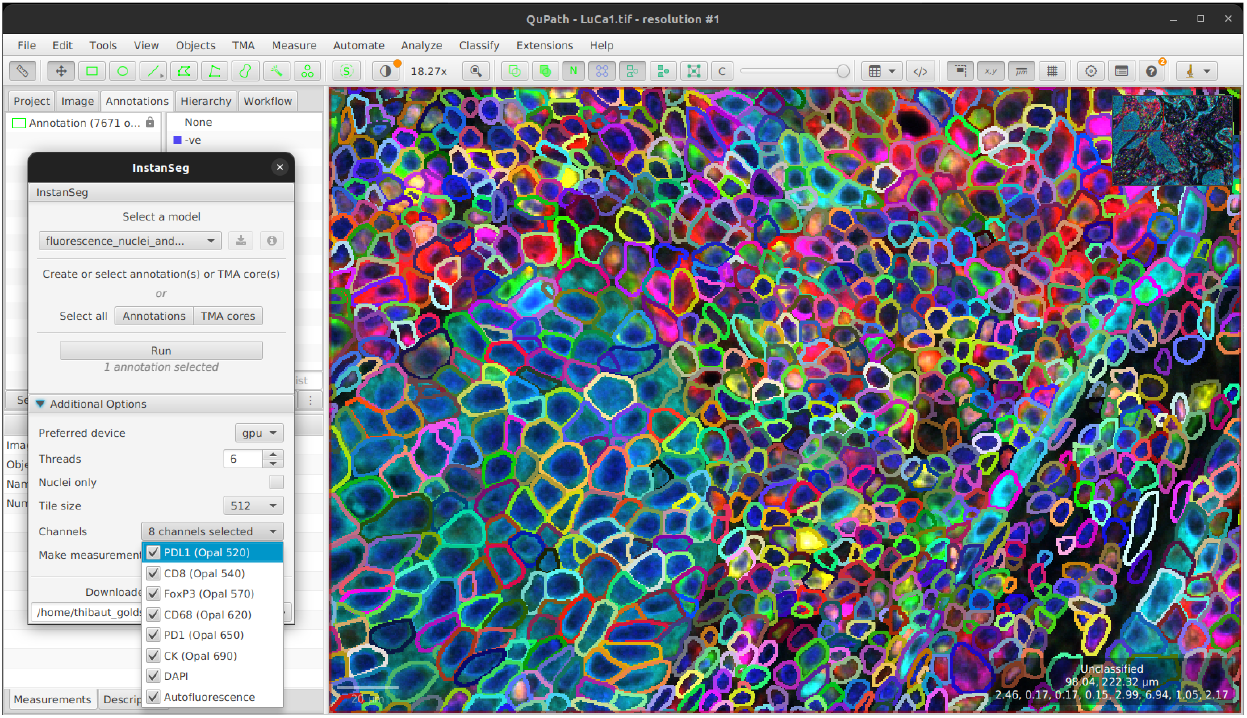
Screenshot showing the interactive InstanSeg extension within QuPath. Our extension provides a user-friendly interface for selecting channels for segmentation, and supports GPU acceleration. Sample multiplexed image from LuCa7 ©Perkin Elmer (CC BY 4.0).

## 4 Discussion

In this work we present a novel method for the simultaneous segmentation of cells and nuclei in multiplexed images. First, we show that InstanSeg improves on the previousstate-of-the-art Mesmer for both nucleus and cell segmentation on the large TissueNet dataset. Not only was the Mesmer method conceptualized using this dataset, a number of ground truth annotations in the validation and test sets were obtained using a Mesmer model. There is a possibility that this provided an advantage to Mesmer when benchmarking on this dataset, yet it was still outperformed by InstanSeg.

InstanSeg is a highly efficient segmentation algorithm, which we have previously shown to achieve similar or greater accuracy to CellPose [8], StarDist [25] and HoVerNet [26] for nucleus segmentation, while reducing processing time by at least 60% [21]. Here, we retain this efficiency while extending our method to support arbitrary input channels and full cell segmentation. With processing speeds of 42.7 images/second, InstanSeg is nearly ten times faster than Mesmer, which is reported to be among the fastest deep learning-based cell segmentation algorithms [9]. The differences in efficiency are largely due to InstanSeg’s lightweight and GPU accelerated postprocessing step. This eliminated postprocessing as a bottleneck, and contrasts with the strategies employed by CellPose or Mesmer, both of which rely upon a computationally expensive flow tracking or a watershed transform to calculate final outputs.

The proposed combination of ChannelNet + InstanSeg enables the segmentation of cells in multiplexed images irrespective of the number and ordering of input biomarkers. This allows the method to generalize across imaging platforms and experiments. Our channel aggregation strategy not only leads to increased segmentation accuracy, but also eliminates the need for users to manually determine the optimal membrane/cytoplasmic markers for the segmentation step making it uniquely easy to apply to new image sets. While histoCAT [18] already enables the prediction of three-channel intermediate representations of multiplexed images, the method requires manual retraining of an Ilastik pixel classifier for different channel combinations. Unlike InstanSeg, this approach risks introducing user-to-user variability and slows down the segmentation workflow. By incorporating additional input channels for segmentation, InstanSeg can more accurately resolve cellular boundaries between biomarkers. We expect that this will allow for improved feature extraction and phenotyping of cells in multiplexed images. Better quantification of cellular properties allows for improved downstream analyses, such as the study of cell interactions within their microenvironments.

Openness and accessibility are central to this work. Our development of ChannelNet benefited from the creators of CPDMI 2023 making a large and heterogeneous multiplexed dataset of hand annotated cells and nuclei freely available [24]. Good training and validation data are crucial, and to our knowledge this is the first such dataset to be shared under a permissive open license. By making our own code freely-available and open-source, and by providing a model pre-trained on CPDMI through a user-friendly QuPath extension, we anticipate that ChannelNet + InstanSeg will provide a new standard baseline for multiplexed cell segmentation in the research community. As the method is applied independently to a wider range of images – acquired using different technologies, to look at even more tissues and markers – we expect that this real-world validation will quickly identify areas where improvement is still required. Our hope is that this will help further accelerate progress by focusing community effort on unsolved problems, and inspire the sharing of new open datasets to train and validate future models.

In conclusion, we have proposed a nucleus and cell segmentation method for multiplexed images using a novel channel invariant module. Our methodology demonstrated substantial improvements in accuracy and efficiency on public segmentation datasets. The combination of ChannelNet + InstanSeg allows for inference on multiplexed images where the number and ordering of channels need not match that encountered during training. Our implementation does not require retraining or any manual intervention, thereby reducing inter-user variability and simplifying segmentation workflows. By providing open-source implementations for both Python and QuPath, in addition to pre-trained models, we have provided tools for biologists to efficiently integrate our method in full analysis pipelines.

## Acknowledgments

We thank Ben Philps and Dr. Pau Carrillo Barberà for their insightful discussions and suggestions that improved this work. We extend our gratitude to the curators of publicly available segmentation datasets used in this work.

TG was supported by the United Kingdom Research and Innovation (grant EP/S02431X/1), UKRI Centre for Doctoral Training in Biomedical AI at the University of Edinburgh, School of Informatics. HB was supported by the EPSRC Visual AI grant EP/T028572/1. This project has been made possible in part by grant number 2021-237595 from the Chan Zuckerberg Initiative DAF, an advised fund of Silicon Valley Community Foundation. This research was funded in part by the Wellcome Trust 223750/Z/21/Z. For the purpose of open access, the author has applied a Creative Commons Attribution (CC BY) licence to any Author Accepted Manuscript version arising from this submission.

---

**Supplementary Fig. 1:**
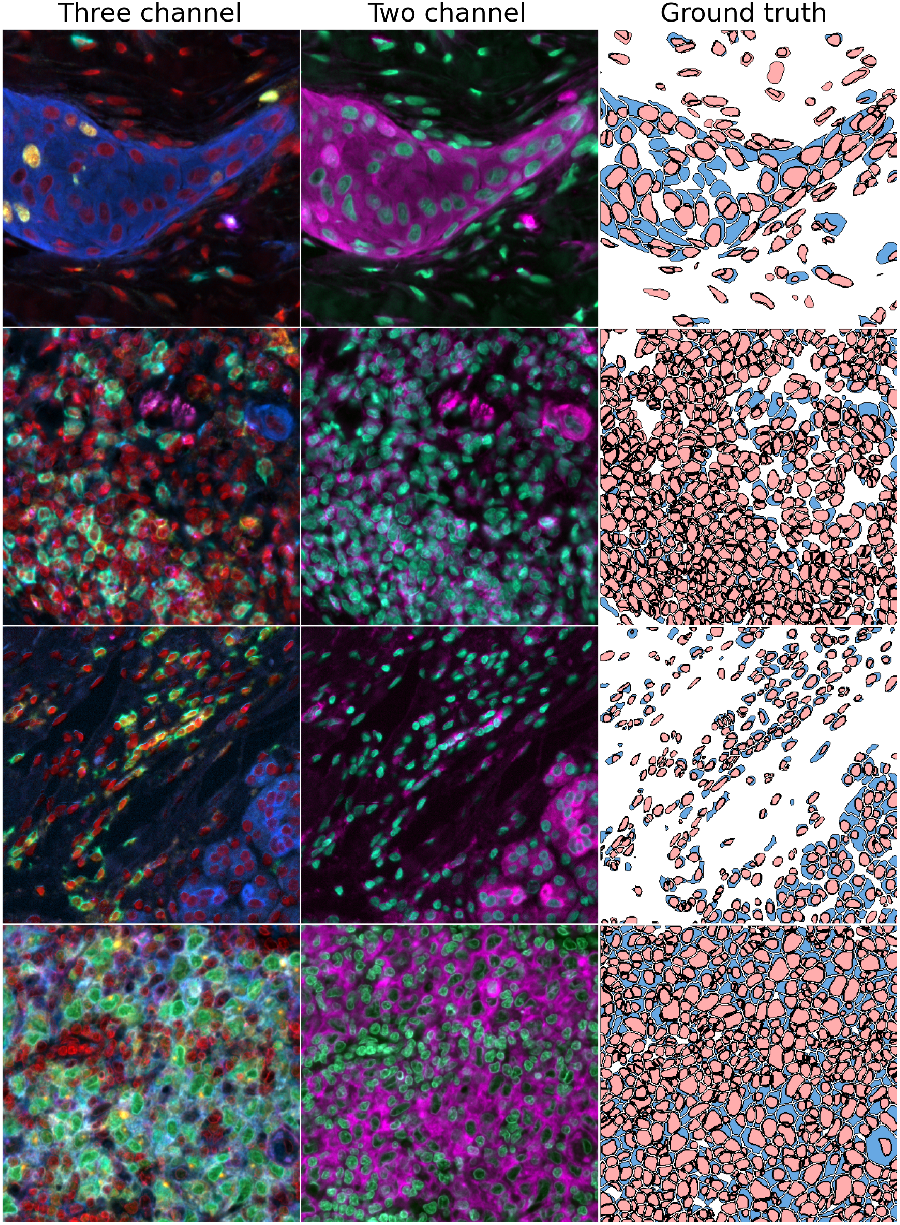
Channel aggregation baselines. Three channel: nuclear biomarkers are kept in a separate channel (here in red) and the other channels are linear projections into RGB space. This is often used in visualisation software. Two channel: markers are separated whether they are mostly expressed in the nucleus or in the cytoplasm, this has previously been used to segment multiplexed images using Mesmer.

**Supplementary Fig. 2:**
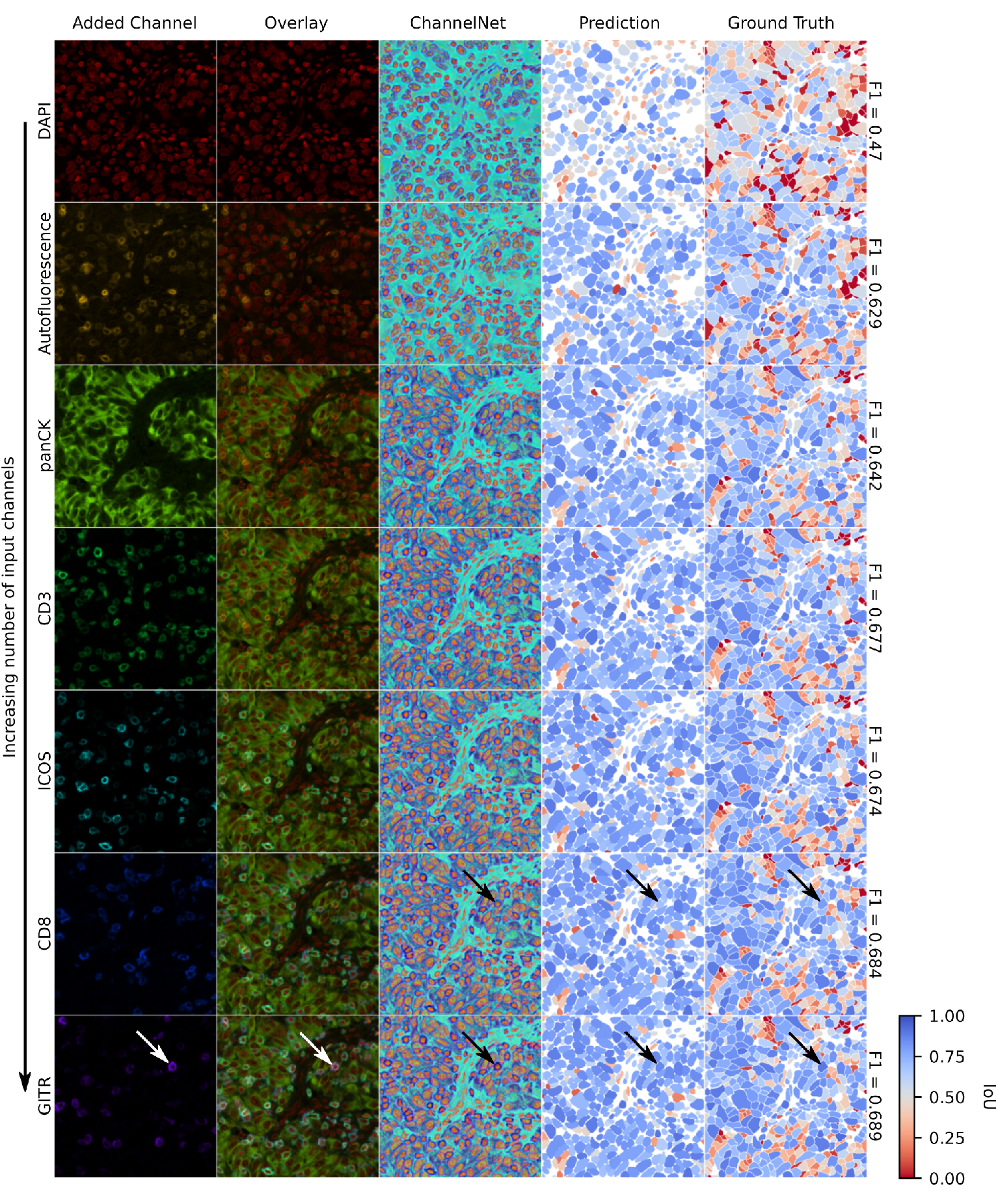
Qualitative results showing the effect of increasing the number of input channels from one (top row) to seven (bottom row). The ChannelNet intermediate RGB representations, which serve as input to the main InstanSeg model are depicted in the central column. The last two columns show the predicted and ground truth whole cell labels, the per-cell agreement is shown using an Intersection over Union (IoU) metric. In other words, instances coloured in red in the columns “Prediction” and “Ground Truth” are false positives and false negatives respectively. Note how markers that were expressed in only some of the cells (e.g. GITR) subtly affected the intermediate RGB representations and eventually allowed for more accurate cellular boundary predictions (see arrows).

## Footnotes

1 https://github.com/stardist/stardist

